# Antimicrobial copper as an effective and practical deterrent to surface transmission of SARS-CoV-2

**DOI:** 10.1101/2022.08.25.505365

**Authors:** Jorge Vera-Otarola, Nicolas Mendez, Constanza Martínez-Valdebenito, Rodolfo Mannheim, Steve Rhodes, Kenneth Lu

## Abstract

The aerosols are critical for SARS-CoV-2 transmission, however in areas with high confluence of people the contaminated surfaces take an important role that we could attack using antimicrobial surfaces including copper. In this study, we wanted to challenge infectious SARS-CoV-2 with two samples of copper surfaces and one plastic surface as control at different direct times contact. To evaluate and quantify virucidal activity of copper against SARS-CoV-2, two methods of experimental infection were performed, TCID_50_ and plaque assays on VeroE6 cells, showing significant inactivation of high titer of SARS-CoV-2 within minutes reaching 99.9 % of inactivation of infectivity on both copper surfaces. Daily high demand surfaces contamination is an issue that we have to worry about not only during the actual pandemic time but also for future, where copper or its alloys will have a pivotal role.

**Importance:** Quantitative data obtained of TCID50 and plaque assay with infectious SARS-CoV-2 virus showed that after direct contact with copper or copper alloys, viruses were inactivated within minutes. Notably, the SARS-CoV-2 virus used in these assays was in high titer (106 PFU/mL) showing strong copper inactivation of the infectious SARS-CoV-2.

## Introduction

The world has been ravaged by Covid-19, with devastating impact on life and economy^1,2^. Significant progress in knowledge and treatment have been made at a break-neck pace, but much work remains to be done in containment and prevention of current and future pandemics. One area of need is the lack of an effective and long-term method for reducing surface transmission of diseases. Antimicrobial detergents effectively kill bacteria and virus quickly, but lack longevity. Industrial nanoparticle coating is expensive and requires disassembly and reassembly of existing fixtures^3^. Spray-on nanoparticle solutions require drying time and though nominally longer-lasting than Lysol, is still not permanent^4^.

Antimicrobial copper has received resurgent attention as a potential solution^5,6^. It received approval by EPA as antimicrobial in 2008^7^, and anti-Covid in February of 2021^8^. With copper being a safe, natural, and ubiquitous compound, it would seem the perfect panacea, however the time to killing of coronavirus is a potential concern. The National Institute of Allergy and Infectious Diseases (NIAID) published data of copper’s superiority in killing coronavirus to plastic and stainless steel surfaces, which essentially had no impact in April of 2020^9^. However, the reported 4 hours required for complete eradication by copper is at odds with other studies on human coronavirus and bacteria, which report significant inactivation on the order of minutes^10–13^. Copper’s mechanism of killing involves electron and oxygen free radicals^7^ causing breakdown of membranes and disintegration of genetic material, thus theoretically work better against simple organisms like virus. If plenty of published reports support copper’s killing time of numerous bacteria in minutes^14–16^, it stands to reason that it would work at least as quickly against viruses. At least one article published subsequent to NIAID study has validated copper’s ability to deactivate SARS-CoV-2 within minutes.^17^ This article by Dr. Keevil also explained the difference in methodology that led to the much longer kill time in the previous paper by NIAID.

The present study seeks to quantify copper’s speed of action against SARS-CoV-2 and shed some light on the practical use of copper as a means to reduce surface transmission. An antiviral activity study comparing 2 different samples of 99.9% copper against plastic surfaces, based on ISO 21702-2019 standard of “Measurement of Antiviral Activity on Plastics and Other Non-porous Surfaces” was performed.

## Material and Methods

### Cell line and SARS-CoV-2 virus isolation

Vero E6 (Vero C1008; ATCC CRL 1586) was grown in Dulbecco’s modified Eagle’s medium high glucose plus L-Glutamine (DMEM; HyClone, Logan, UT) containing 10% fetal bovine serum (HyClone, Logan, UT, USA), 1% amphotericin B, 1% penicillin-streptomycin and at 37°C in a 5% CO_2_ atmosphere. Infectious virus SARS-CoV-2 was isolated from a Chilean’s patient during pandemic period on June 2020 and incubated on Vero E6 cells at Biosafety Level 3 laboratory. Briefly, 200 µL nasopharyngeal swab sample from a SARS-CoV-2 positive RT-qPCR patient was incubated with 400 µL de solution B (DMEM media with 50 µg/mL Gentamicin, 5 µg/mL amphotericin B) for 30 minutes at room temperature. Treated sample was incubated on Vero E6 cells for 1hour at 37ºC and sample was replaced by Dulbecco’s modified Eagle’s medium high glucose plus Glutamax (# 10566-016, Gibco BRL, Life Technologies Corporation, Carlsbad, CA, USA) containing 2% fetal bovine serum (FBS, HyClone, Logan, UT, USA), 10mM HEPES (#25-060-Cl, Corning), 1% amphotericin B (Ampho B, #30-003-CF, Corning), 1% penicillin-streptomycin (P/S, #30-002-Cl, Corning), 1% nonessential amino acids (#25-025-Cl, Corning) at 37°C in a 5% CO_2_ atmosphere. During the days post-infection, the supernatants were monitored using Standard Q COVID-19 Antigen Test (SD Biosensor Inc, Korea) until positive signal was captured from the assays using mock-infected cells as negative control. Viral stock was confirmed by RT-qPCR (LightMix® SARS-CoV-2 RdRp-gene EAV PSR & Ctrl (TIB MOLBIOL). Reactions were run in a LightCycler® 480 real time-PCR system (Roche). Viral titers were obtained by plaque assay.

### Copper test samples

Two samples of copper foil, both >99.9% copper content, were used in this experiment. One is a copper (certified 99,99% UNS C 12200 Cu DHP) sheet with adhesive applied to the underside (The Clean Copper Company, Los Angeles, CA), hereafter called Copper Sticker, and the other (99.93%), without adhesive, hereafter called Copper Coupon. Materials are shown in figure 1 and 4.

**Figure 1.**
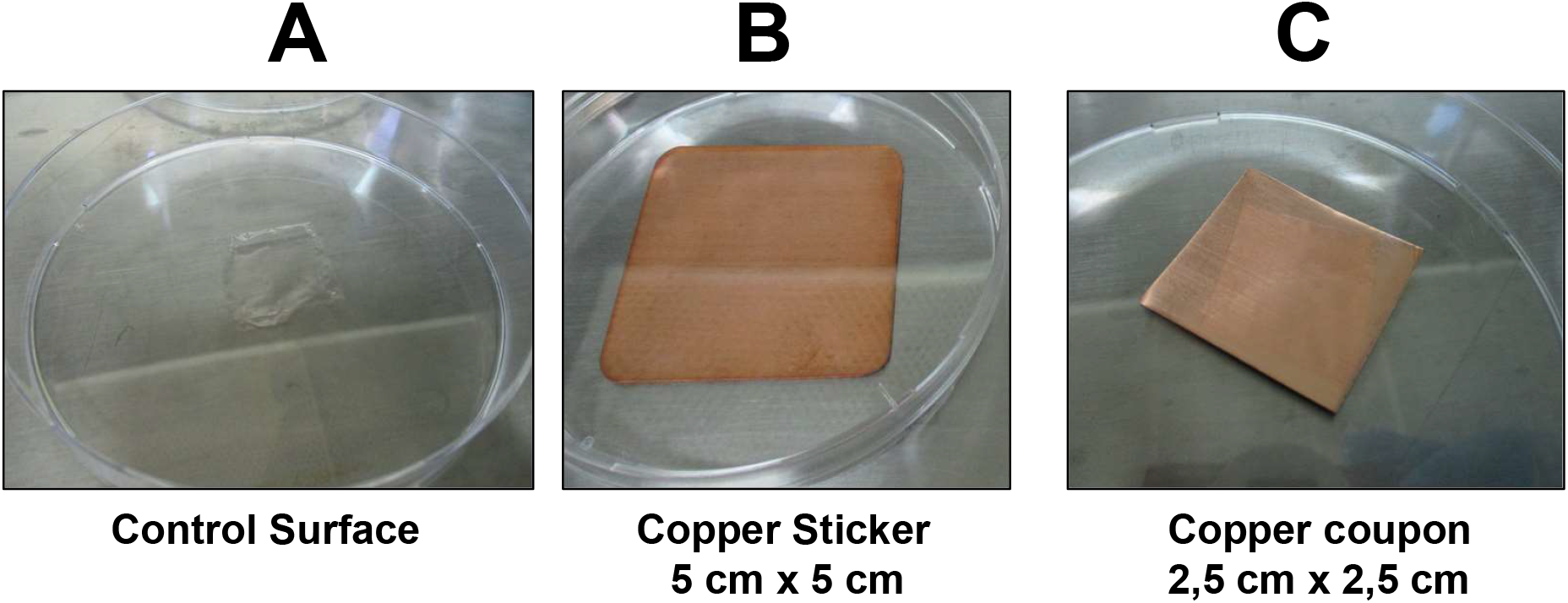
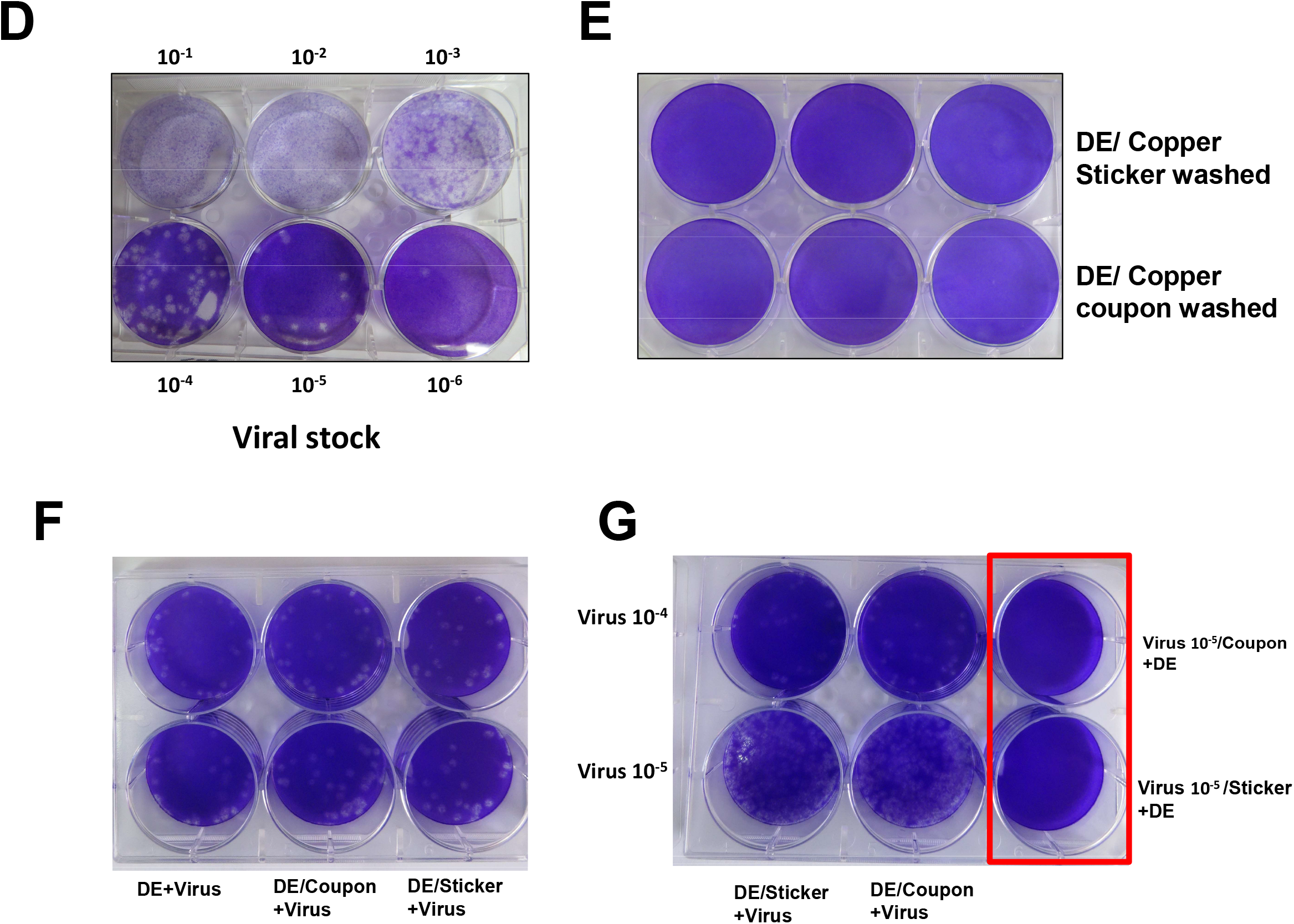
Surface format for virucidal activity measurements. **A)** Control surface: Cell Culture Petri dish, **B)** Copper sticker, **C)** Copper coupon. **D)**Viral stock used in this study measured by plaque assay in VeroE6 cells described in methods. **E)** Cytotoxicity assay of surfaces plus neutralizer DE. **F)** Representative plaque assay for Susceptibility assay described in methods. **G)** Representative plaque assay for neutralization assay as described in methods. Two viral titer were added independently to washed copper surface solution to the two upper and bottom wells showing neutralization. The last upper and bottom well (red square), one titer of virus was incubated with copper surface for 1 hour and neutralizer DE was added stopping virucidal activity of copper surfaces.

### Virucidal Activity Measurement by Plaque Assay and TCID_50_ Assay

Experiments based on standard ISO 21702-2019 “Measurement of antiviral activity on plastics and other non-porous surfaces” were performed with copper sticker and coupon. Each copper sample was placed inside 10 cm cell culture plate. 100 microliters of virus SARS-CoV-2 (viral titer 4.6×10^6^ PFU/mL) was placed in contact with copper or directly on culture plate as negative surface control. The droplet of virus was covered by 400 mm^2^ of propylene film, spreading the virus evenly throughout the film, making sure that virus inoculum does not leak beyond the edges of the film. After predesignated contact times, 2.5 mL of Dey-Engley (DE) neutralizer (# D3435, Sigma Aldrich) was used to recover viruses from both propylene and test surfaces. 10-fold dilutions were prepared in EMEM 1X ((EMEM Lonza, #12-684F), 0,12% NaHCO_3_ (Sigma, S5761), 10mM HEPES, 1X L-glutamine (#25-005-Cl, Corning), 1% amphotericin B, 1% penicillin-streptomycin. Virus dilutions were added to each well (250 µL of volume) with confluent monolayer of Vero E6 cells for 1 hour at 37°C in a 5% CO_2_ atmosphere with continuous slanting of the plate by a rotator. Virus was removed and 2 mL of overlay media (EMEM 1X Lonza 12-684F, 5 % FBS HyClone, 0.6 % purified Agar OXOID #LP0028, 0.12% NaHCO3 Sigma #S5761, 0.01% Dextran Sigma #D9885) was added. 3 days post-infection 4 mL of paraformaldehyde 4% (#158127, Sigma) was added and incubated overnight at 4ºC. The semisolid agar was removed and 0.5% crystal violet was added to each well for 2 minutes. The plates were washed of crystal violet, dried and the plaques were counted for viral titer and registered manually in PFU/ mL. The log of reduction for plaque assay was calculated as (mean of Log PFU/ mL of control) - (mean of Log PFU/ mL at each time contact). The percentage of reduction was calculated using the formula % reduction = (1-10^ - (log of reduction) x 100). In order to show control surface had no effect on the virus, the PFU values with exposure to control surfaces at 30 seconds and 60 minutes were compared by this formula: [(Log maximun number of PFU - Log minimun number of PFU) / Log mean of PFU]. A value of ≤ 0.2 indicates no difference.

In parallel, TCID_50_ assays were also used to calculate the reduction of infectious SARS-CoV-2 virus. Same 10-fold dilutions used for plaque assays were used to infect Vero E6 cells in 96-well plates using 100 uL of each dilution. 3 days post-infection the cytopathic effect was recorded by microscopy and later the plates were stained with crystal violet as above. The unstained cells (Lysis) in each well were considered infected cells, and stained cells as uninfected cells. The data was analysed using the Spearman Karber method [Negative Log of TCID_50_/100 µL = −log of 1st dilution assayed – [((Σ of % mortality at each dilution/100) – 0.5) x (log of dilution)]. The log of reduction was calculated as (LogTCID_50_/control) - (LogTCID_50_/ each time contact). The percentage of reduction was calculated using the formula % reduction= (1-10^ - (log of reduction) x 100). All infection assays were performed in duplicate or triplicate plates at least in two independent experiments. Statistical analysis was performed using ANOVA and Kruskal-Wallis multiple comparison test (P **<** 0.05).

### Cytotoxicity, Susceptibility Verification and Neutralization

Absence of sample and control surface cytotoxicity on Vero E6 cell line was verified by passing Dey Engley neutralizer over copper and control surfaces, then incubating on Vero E6 cells for 1 hour. Overlay media was added and 3 days post-infection 3 mL of paraformaldehyde 4% was added and incubated overnight at 4ºC. The semisolid agar was removed and 0.5% crystal violet was added to each well for 5 minutes. Staining indicates survivability of Vero E6 cells. Susceptibility of Vero E6 cells to SARS-CoV-2 was tested by washing the sample and control surfaces with 2.5 mL of Dey-Engley neutralizer (DE). Two mililiters of washed was recovered in a 15 mL tube, where 50 µL of SARS-COV-2 virus with a viral titer of 5×10^4^ PFU/mL were added and incubated 30 minutes at room temperature. A volume of 250 µL of each tube was inoculated on wells with a monolayer of Vero E6 cells and plaque assay as above was performed. This is compared to Dey-Engley neutralizer added directly to the virus without prior exposure to sample and control surfaces. Similar procedure is performed in order to evaluate neutralization of virucidal activity of copper surfaces. The surfaces are washed with 2.5 mL of DE and the washed recovered is incubated with 100 µL of SARS-COV-2 virus with a viral titer of 5×10^4^ or of 5×10^5^ PFU/mL, incubated for 1 hour and 250 µL of mix is added to cells. In parallel, 100 µL of SARS-COV-2 virus with a viral titer of 5×10^5^ is incubate on copper surface for 1 hour, 2.5 mL of neutralizer DE is added to virus/surface and recovered in a 15 mL tube, where 250 µL of mix is added to cells. Plaque assays were performed with those samples as above.

## Results

The 2 sample surfaces and the control surface tested are shown in Figure 1. Figure 1A shows the control surface, which is a cell culture petri dish. The copper sticker is shown in Figure 1B, and the copper coupon is shown in Figure 1C. SARS-CoV-2 viral stock titration is shown in Figure 1D. Control experiment shows in Figure 1E, that both copper surfaces were not cytotoxic to Vero E6 cells. Susceptibility of Vero E6 cells to virus is shown in Figure 1F, where it was the same between virus/neutralizer alone and virus/surface washed neutralizer. The PFU value for virus/neutralizer alone (control) was 3.0×10^3^ PFU/ mL. The PFU value for virus/copper sticker-washed neutralizer was 2.92×10^3^ PFU/mL and the PFU value for virus/ copper coupon-washed neutralizer was 2.95×10^3^. These results shows that both copper surfaces are not altering susceptibility of VeroE6 cells to virus. Regarding to neutralization of virucidal activity, as is shown in Figure 1G, the neutralizer DE supress virucidal activity from copper coupon and copper sticker using two different viral titer. Notably, at 1 hour of exposure of virus (titer 5×10^5^ PFU/mL) with both copper surfaces, there is not plaque visualized (red square). These preliminary results suggest complete inhibition of SARS-CoV-2 infectivity by both copper surfaces in 1 hour of exposition.

The virucidal activity was measured based on standard ISO 21702-2019, where 100 uL of SARS-CoV-2 infectious virus (titer 4.6×10^6^) was placed on surface and incubated for 30 seconds, 60 seconds, 5, 15, 30 and 60 minutes, covered by a 400 mm^2^ propylene film. The results show that the virus recovered from surface control at 30 seconds and 60 minutes yielded a reproducibility value of 0.015, validating the control surfaces (Figure 2A, 2B; Table 1 and 2). The formula used is (*Lmax*-*Lmin*)/ (*Lmean*) ≤ 0,2, where *Lmax* is the common logarithm (i.e. base 10 logarithm) of the maximum number of plaques recovered from a control Surface. *Lmin* is the common logarithm of the minimum number of plaques recovered from a control surface, and *Lmean* is the common logarithm of the mean number of plaques recovered from the control surfaces.

**Table 1.** Values of virucidal Activity on Copper’s sticker compared to control surface.

The PFU/mL was calculated following the formula PFU/mL= (N° of plaque x reciprocal value of dilution)/ mL added to well. The log TCID<sub>50</sub>/100µL was calculate using the Spearman-Karber method. Experiments were done in triplicate plates at least in two independent experiments.
| **Surface** | **Time of contact** | **PFU/mL** | **Log TCID<sub>50</sub>/0.1mL** | **% Reduction**<br>**Plaque Assay** | **% Reduction**<br>**TCID<sub>50</sub>** |
| --- | --- | --- | --- | --- | --- |
| Control | 30 sec | 9.60 x 10<sup>6</sup> | 4.65 | 0 | 0 |
| Control | 60 min | 8.32 x 10<sup>6</sup> | 4.02 | 0 | 0 |
| Cu Sticker | 30 sec | 7.84 x 10<sup>6</sup> | 4.15 | 18 | 68.38 |
| Cu Sticker | 60 sec | 7.68 x 10<sup>6</sup> | 4.02 | 20 | 76.56 |
| Cu Sticker | 5 min | 8.48 x 10<sup>5</sup> | 3.27 | 91 | 95.83 |
| Cu Sticker | 15 min | 0 | 0.90 | 100 | 99.98 |
| Cu Sticker | 30 min | 0 | 0.90 | 100 | 99.98 |
| Cu Sticker | 60 min | 0 | 0.90 | 100 | 99.98 |

**Table 2.** Values of virucidal Activity on Copper coupon compared to control surface.

The PFU/mL was calculated following the formula PFU/mL= (N° of plaque x reciprocal value of dilution)/ mL added to well. The log TCID<sub>50</sub>/100µL was calculate using the Spearman-Karber method. Experiments were done in triplicate plates at least in two independent experiments
| **Surface** | **Time of contact** | **PFU/mL** | **Log TCID<sub>50</sub>/0,1mL** | **% Reduction Plaque Assay** | **% Reduction TCID<sub>50</sub>** |
| --- | --- | --- | --- | --- | --- |
| Control | 30 sec | 9.76 x 10<sup>6</sup> | 4.65 | 0 | 0 |
| Control | 60 min | 6.72 x10<sup>6</sup> | 4.58 | 0 | 0 |
| Cu Coupon | 30 sec | 9.6 x 10<sup>6</sup> | 4.52 | 1.64 | 25.01 |
| Cu Coupon | 60 sec | 7.84 x 10<sup>6</sup> | 4.15 | 19.67 | 57.83 |
| Cu Coupon | 5 min | 7.2 x 10<sup>6</sup> | 4.27 | 26.23 | 68.38 |
| Cu Coupon | 15 min | 2.88 x 10<sup>5</sup> | 3.27 | 97.05 | 95.78 |
| Cu Coupon | 30 min | 0 | 1.02 | 100 | 99.98 |
| Cu Coupon | 60 min | 0 | 1.20 | 100 | 99.97 |

**Figure 2.**
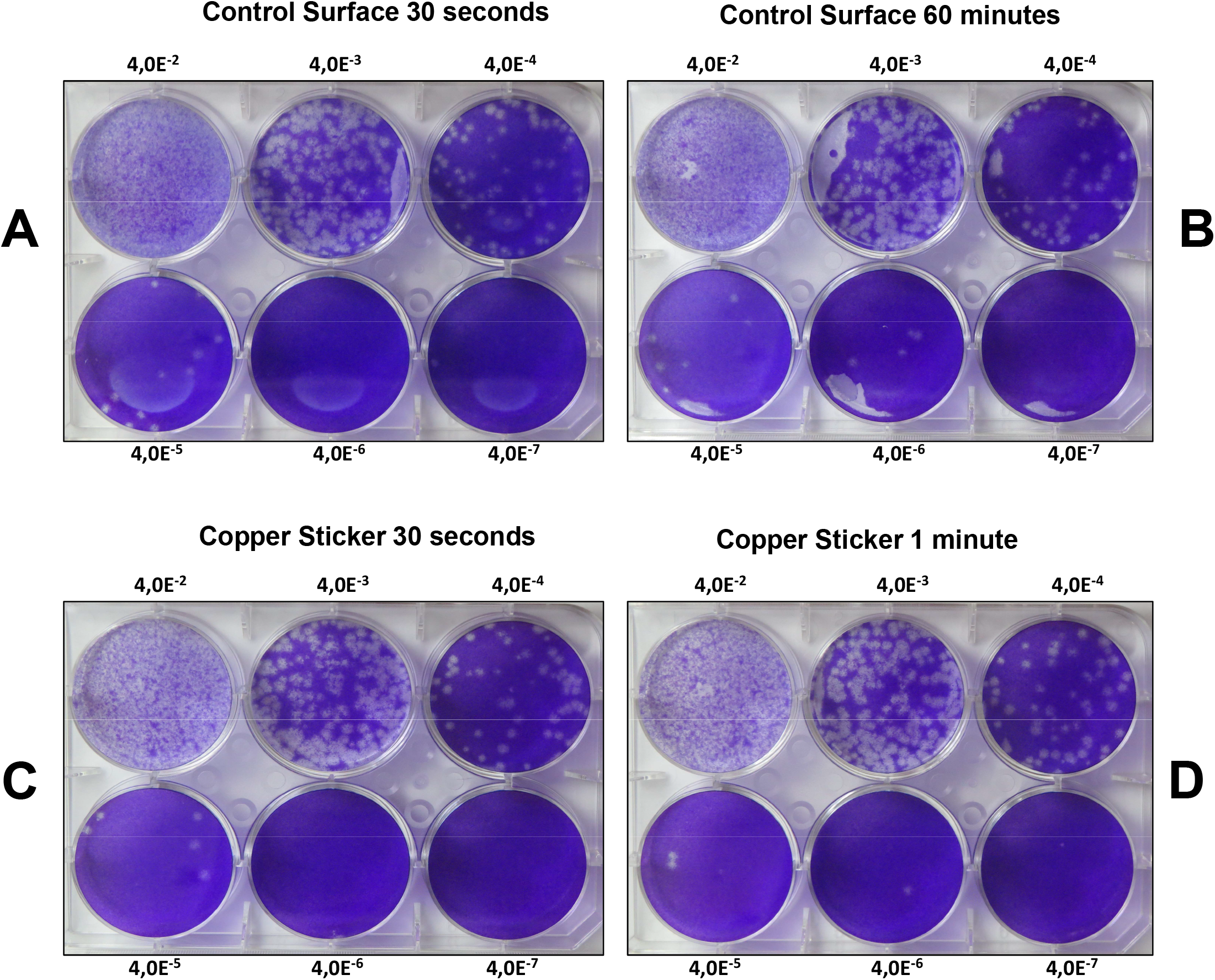
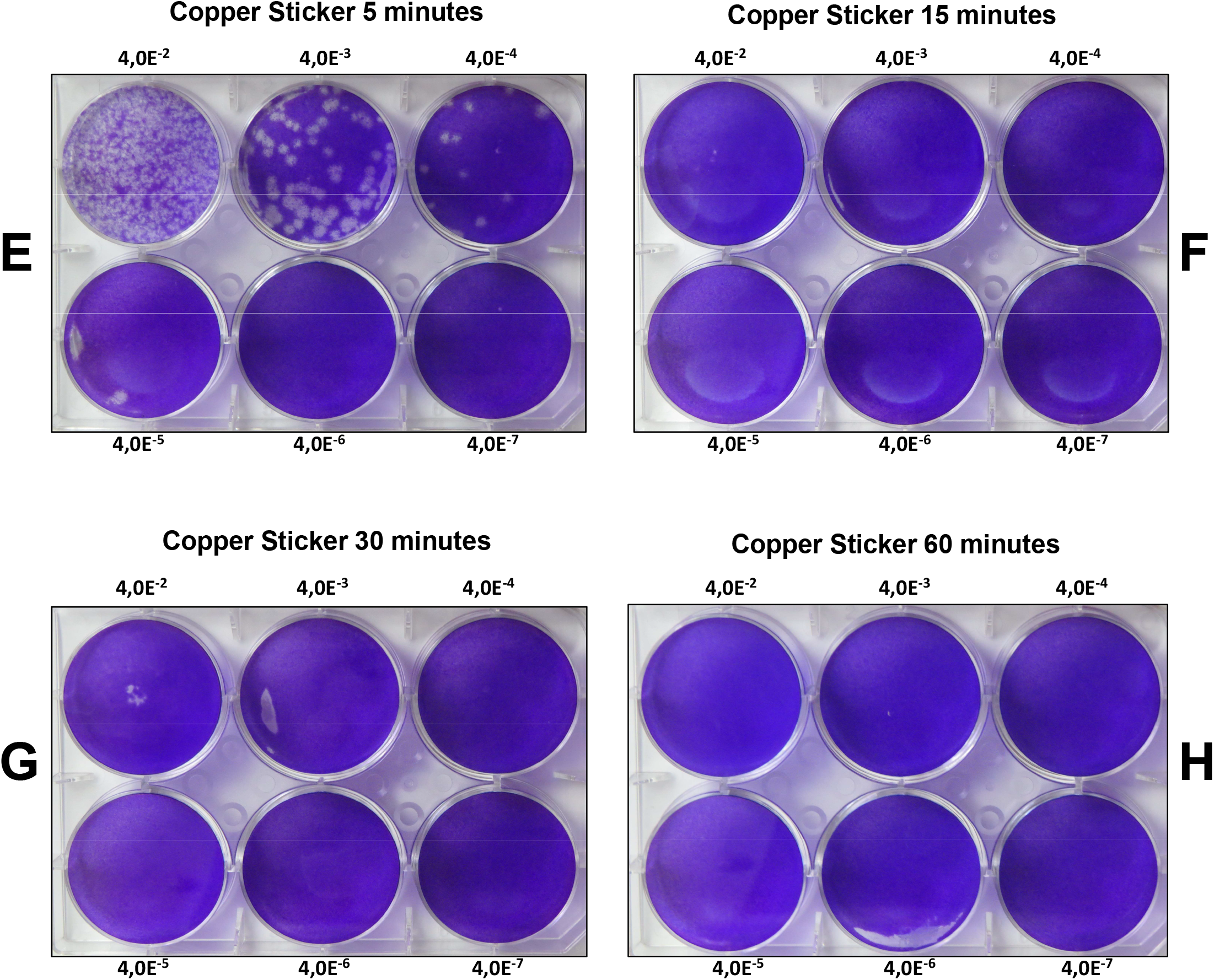
Copper Sticker reduces infectious SARS-CoV-2 virus after defined time contact by plaque assay. Exposition of SARS-CoV-2 virus with copper sticker at different times of contact determined by plaque assay. **A-B)** Control surface at 30 seconds and 60 minutes of contact. **C-H)** Copper Sticker surface at 30, 60 seconds and 5, 15, 30 and 60 minutes of contact.

At 30 and 60 seconds of contact of the virus with copper surfaces there was not a significant reduction of infectious virus (for copper sticker see figures 2C, 2D, 3B; for copper coupon see figures 5C, 5D, 6B; Values in Table 1 and 2). At 5 minutes of contact between virus and copper sticker, 91% of reduction was obtained by plaque assay (figure 2E, values in Table 1) and 95% by TCID_50_ (figure 3C on left, values in Table 1). At same time contact with copper coupon, 26% of reduction was obtained by plaque assay (figure 5E, values in Table 2) and 68% by TCID_50_ (figures 6C on left, values in Table 2). Starting at 15 minutes of contact between copper coupon and SARS-CoV-2 there was 1 log reduction of infectious SARS-CoV-2 equivalent to 97.05 % by plaque assay (figure 5F, values in Table 2) and 95.78% by TCID_50_ assay (figure 6C on right, values in Table 2). Stronger reduction of SARS-CoV-2 infectious virus was obtained with copper sticker at 15 minutes of contact time, where 100% by plaque assay (figure 2F, values in Table 1) and 99.98% by TCID_50_ assay (figure 3C on right, Values in Table 1). When the time contact was higher, such as 30 or 60 minutes, zero plaques were registered on plaque assay, and most of cells were non-infected on TCID_50_ assay (for copper sticker see figures 2G, 2H and 3D; for copper coupon see figures 5G, 5H and 6D; Values in Table 1 and 2). Statistical analysis for all experiments are shown in figure 7, suggesting that copper coupon significantly reduce infectious SARS-CoV-2 virus in less than 30 minutes and copper sticker with Clean Copper technology reduce infectious SARS-CoV-2 virus in less than 15 minutes.

**Figure 3.**
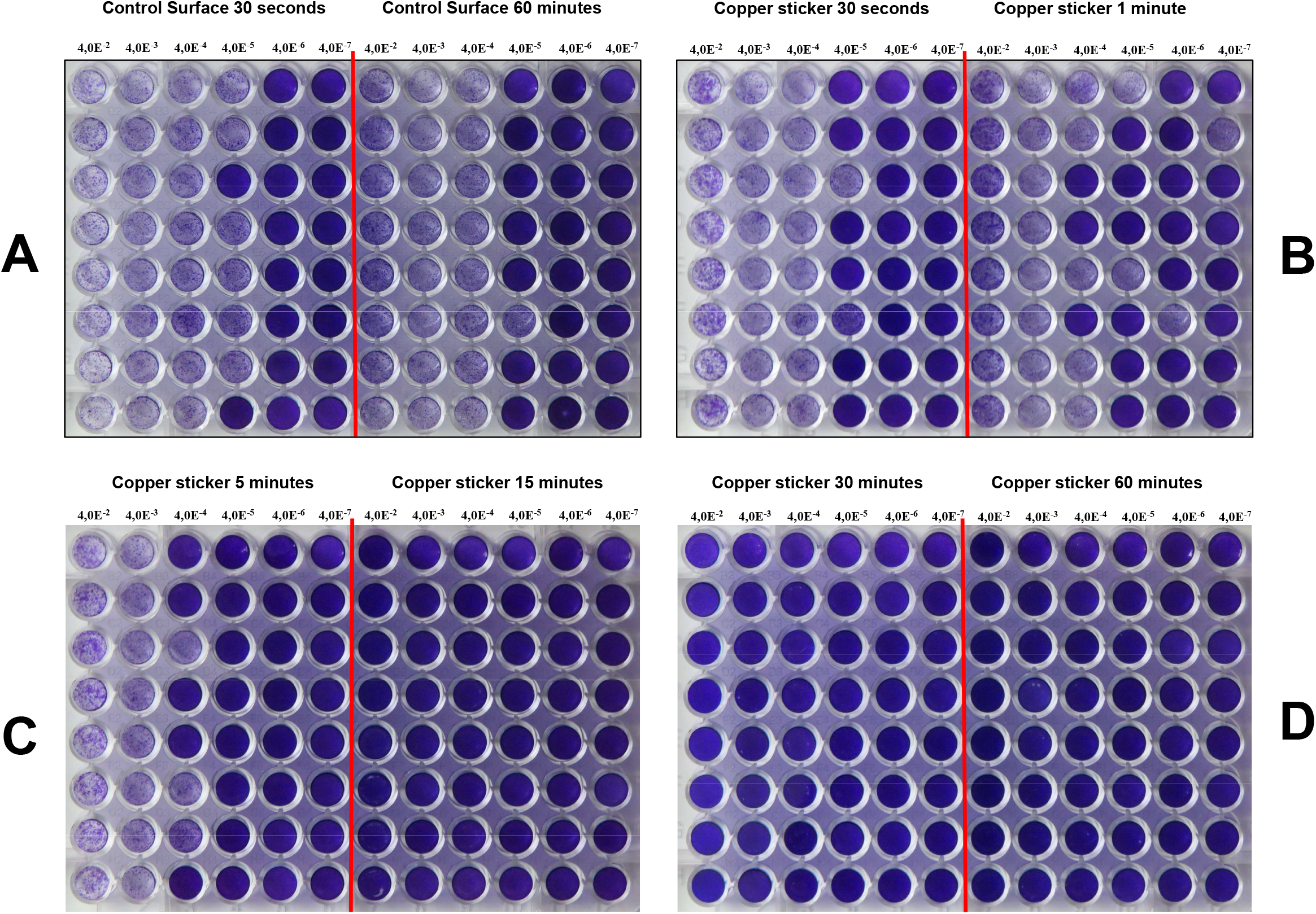
Copper Sticker reduces infectious SARS-CoV-2 virus after defined time contact by TCID_50_ assay. Exposition of SARS-CoV-2 virus with copper sticker at different times of contact determined by TCID_50_ assay. **A)** Control surface at 30 seconds and 60 minutes of contact. **B-D)** Copper Sticker surface at 30, 60 seconds and 5, 15, 30 and 60 minutes of contact.

**Figure 4.**
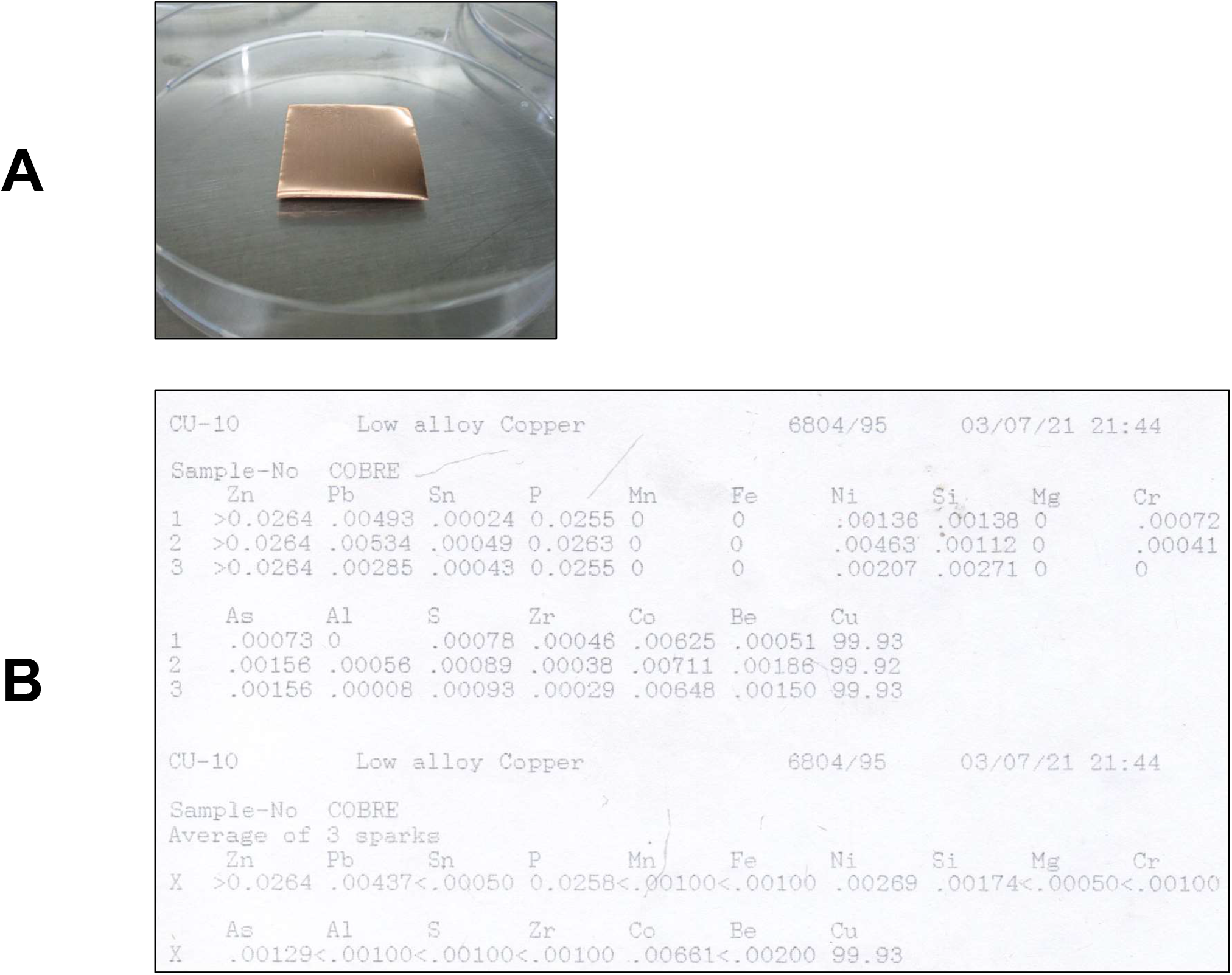
Features of Copper coupon used in this study. **A)** Picture of copper coupon in petri dish plate used for assays. **B)** Optical emission spectrometry analysis of Cu DHP, three sparks and the average of the three trials.

**Figure 5.**
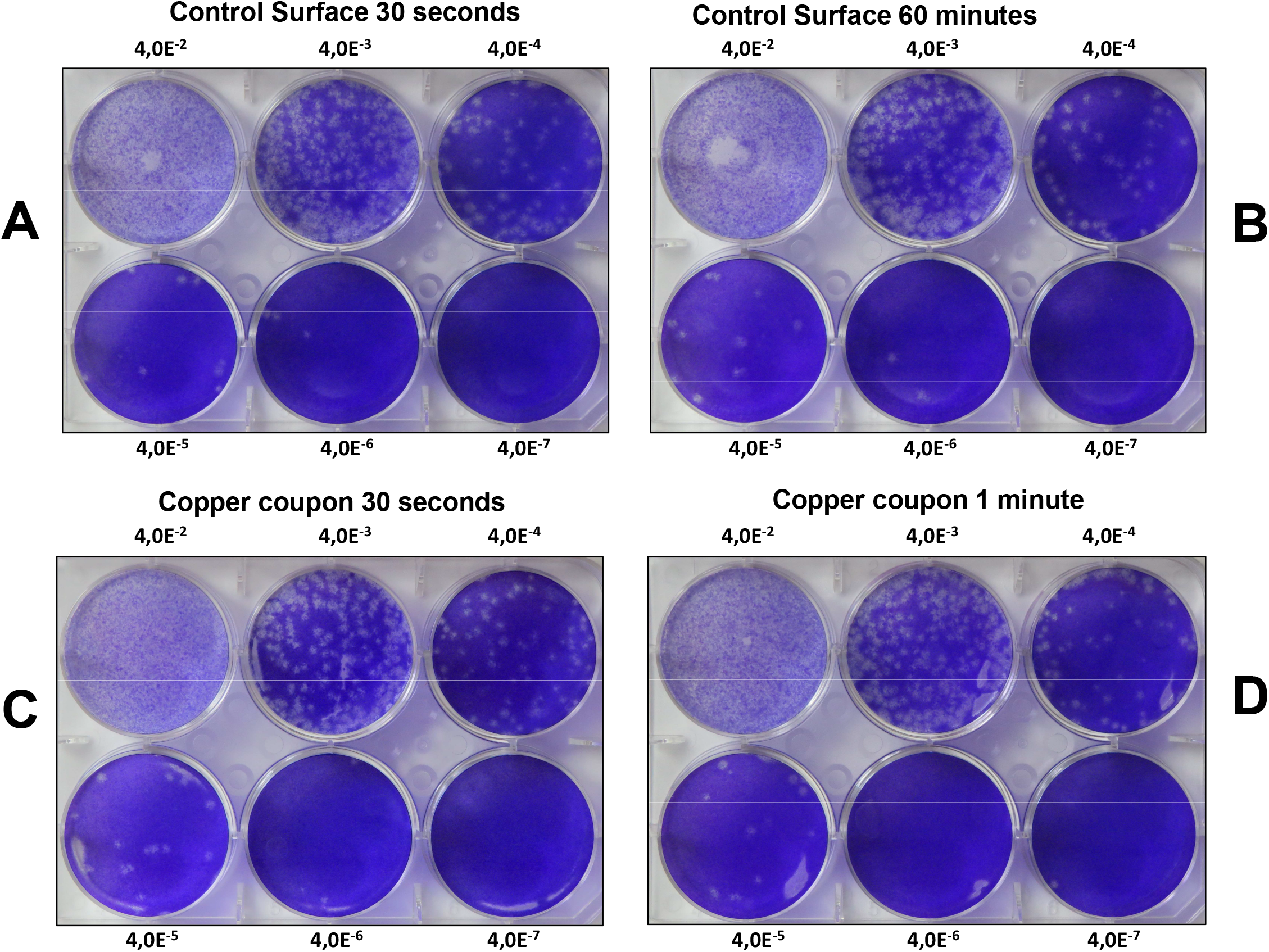
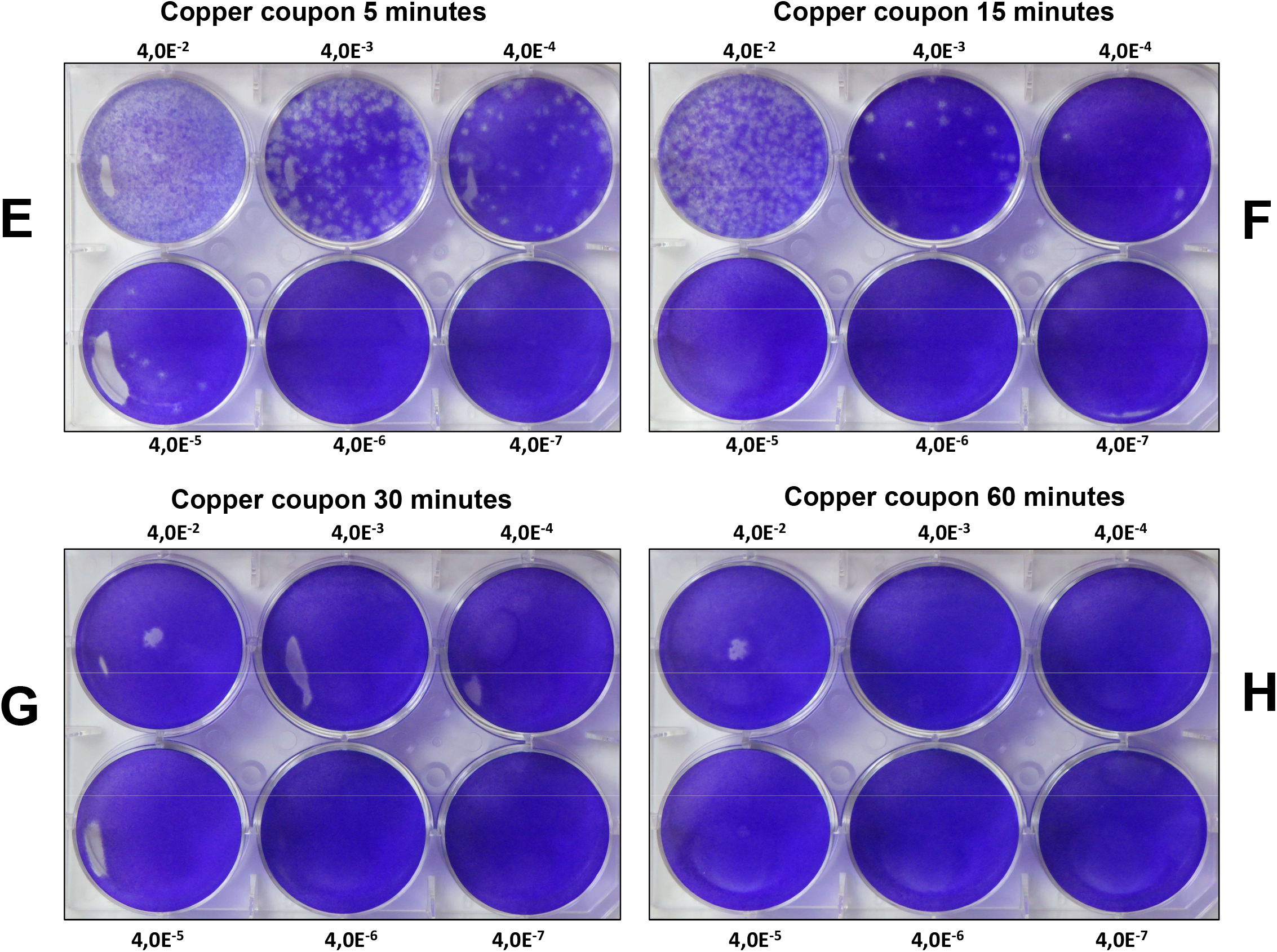
Copper coupon reduces infectious SARS-CoV-2 virus after defined time contact by plaque assay. Exposition of SARS-CoV-2 virus with copper coupon at different times of contact determined by plaque assay. **A-B)** Control surface at 30 seconds and 60 minutes of contact. **C-H)** Copper coupon surface at 30, 60 seconds and 5, 15, 30 and 60 minutes of contact.

**Figure 6.**
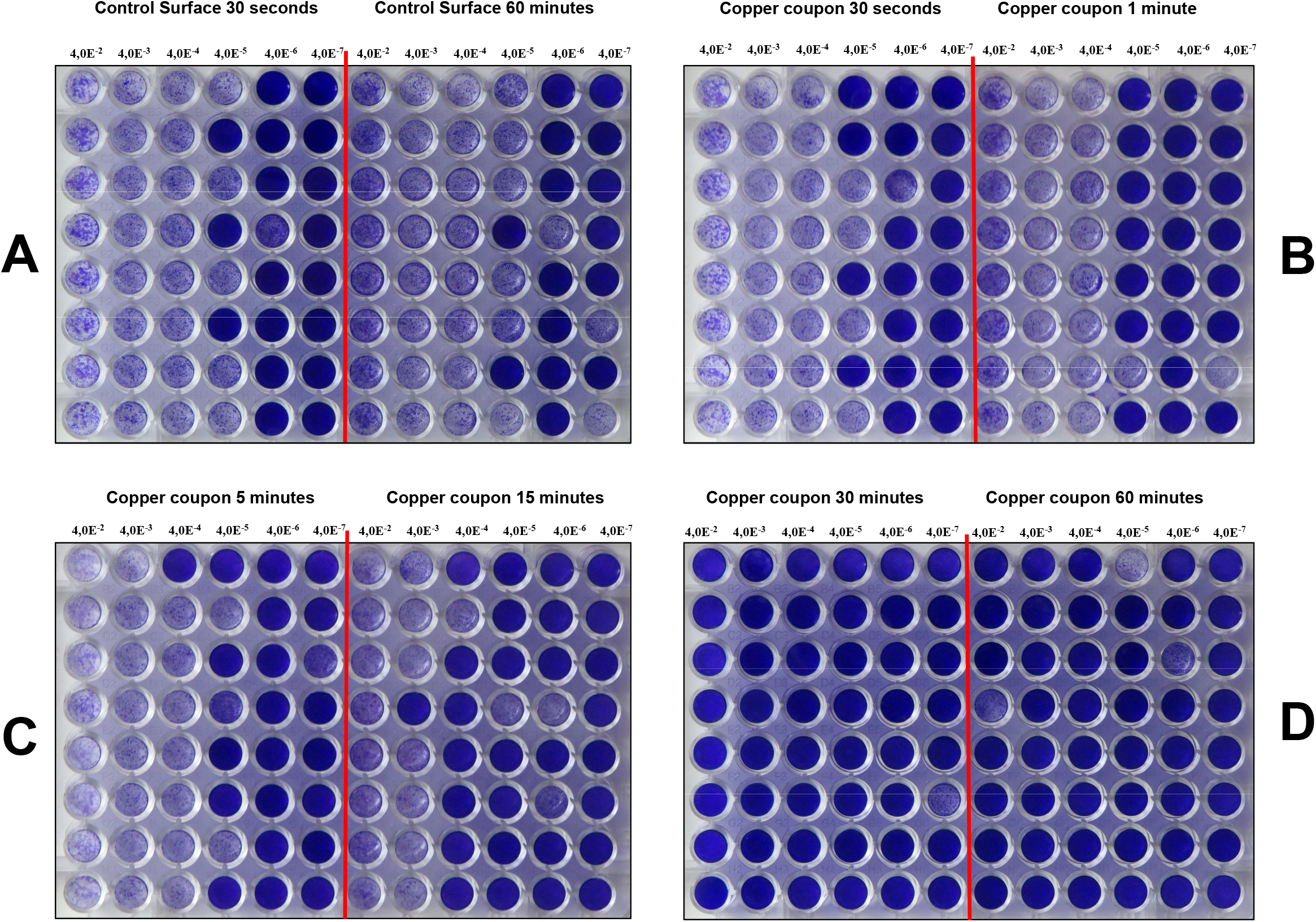
Copper coupon reduces infectious SARS-CoV-2 virus after defined time contact by TCID_50_ assay. Exposition of SARS-CoV-2 virus with copper coupon at different times of contact determined by TCID_50_ assay. **A)** Control surface at 30 seconds and 60 minutes of contact. **B-D)** Copper coupon surface at 30, 60 seconds and 5, 15, 30 and 60 minutes of contact.

**Figure 7.**
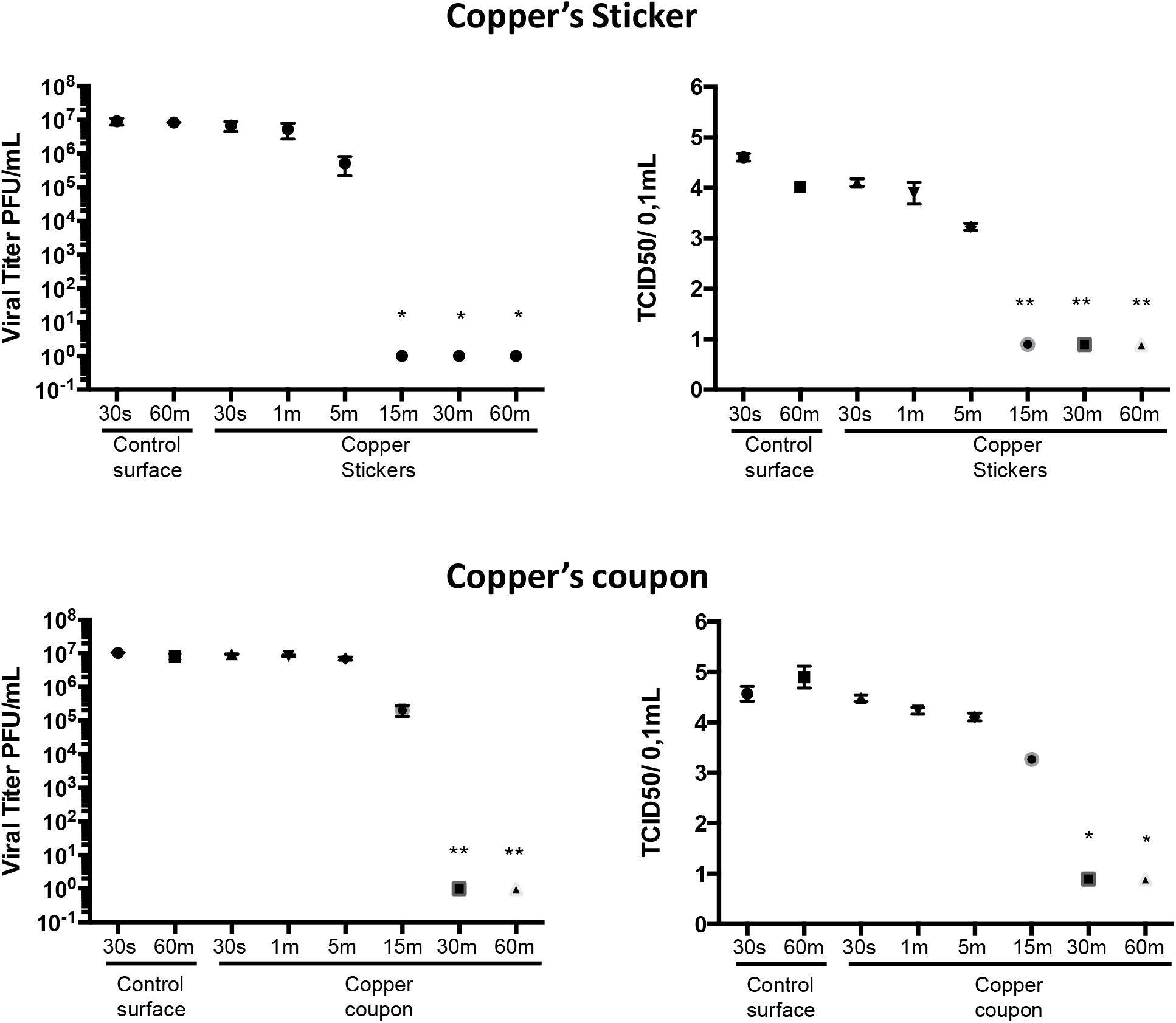
Copper surfaces reduces infectious SARS-CoV-2 virus after defined time contact by plaque assay and TCID_50_. All data from plaque assay and TCID50 were plotted. Values are the mean ±SD of results shown in table 1 and 2. Statistical analysis was performed by the ANOVA test followed by Kruskal-Wallis multiple comparisons, *P < 0.05 versus the control surfaces at 30seconds of contact.

## Discussion

Tests were carried out to evaluate the antiviral activity of copper sheets following the ISO 21702-2019 Standard, by comparing the amount of infectious SARS-COV-2 virus obtained after contact with the copper stickers (Clean Copper, LLC) compared to plastic surfaces, at times of 1, 5, 15, 30 and 60 minutes of contact. The controls required to validate the materials, methodology and the copper sheets used in this study were carried out. Controls assays on figure 1 validate the test, not observing cytotoxicity in the target cells of infection mediated by the copper sheets, nor does the use of the neutralizer in contact with the copper sheets increase or decrease the susceptibility or permissiveness of the cells to infection by the SARS-COV-2 virus. In addition, the neutralizer suppresses the virucidal activity. Figures 2, 3, 5 and 6 show the results of the antiviral activity, where the reduction of infectious virus SARS-COV-2 is calculated after contact with plastic surfaces as a control or on copper sheets.

The results show that the recovery of the infectious SARS-COV-2 virus in contact with the culture plate were similar at all times of the test, which rules out the instability of the virus at the times used in the tests and validates the surface as a control. In the case of the recovery of infectious SARS-COV-2 virus after contact with copper, the results show that there is a steady reduction of infectious virus SARS-COV-2 starting as the earliest time point of 1 minute contact time, achieving 91% on plaque assay and 75.6% on TCID_50_ at 5 minutes, and complete inhibition of infectivity after 15 minutes of contact.

It is expected that the two different types of viral survivability tests offer slight differences, as they examine different aspects of viral activity. TCID_50_ measures via visualization the cytopathic effect in the wells (qualitative), while plaque assay measures actual numbers of plaques (quantitative). Moreover, the viruses have greater mobility to infect neighbor cells in TCID_50_ assay as compared to plaque assay, where overlay media (semi solid) restricts the infection of neighbor cells. The differences appear more prominently early on in incubation, but disappear after 15 minutes of contact, as complete eradication is seen.

In the published report by the NIAID^9^, even though 4 hours was the reported time at which no survivable SARS-CoV-2 was recovered, it also showed an exponential decay in viral survivability starting early. Also, the timepoints of sampling were not reported, instead the decay rates were estimated using a Bayesian regression model. It gives a more qualitative assessment of survivability in aerosol and other surfaces with estimated time decay model and half-lives. In the present study, actual viral titers were measured at specific timepoints, giving a more accurate timeline of viral decay.

These results show that copper sticker and copper coupon have virucidal activity against SARS-CoV-2 and results are in line with results obtained by Keevil’s lab^14,^ where they show that copper has virucidal activity in minutes after contact of coronavirus with copper. Notably, the reduction showed in this study is using a very high viral titer (10^6^ PFU/mL) which is difficult to find naturally in the environment, further reinforcing the idea that copper is a potent virucidal agent against SARS-CoV-2.

Of note, respiratory droplet transmission is likely the predominant route of transmission, however fomite transmission remains an important route. In an epidemiologic and tracing study by the Institute of Public Health, Guangzhou Medical University, 2 family clusters under strict movement monitoring were shown to have most likely been initially infected via touch of contaminated elevator buttons^18^. Both the WHO and CDC have emphasized the risk of fomite transmission of Covid-19. A detailed viral sequencing and tracing study on a Covid-19 outbreak in St. Augustine Hospital in Durban, South Africa in March and April of 2020 pointed to fomite transmission as the predominant route^19^. After masking, environmental disinfection remains an import means of contagion prevention.

These results show that copper can be a practical and effective deterrent to surface transmission of Covid-19. Its onset of action is quick, and its effect is permanent, protecting surfaces against coronavirus on a perpetual basis. Additionally, copper’s proven ability to reduce hospital acquired infection (HAI)^20^ and its efficacy against essentially all microbes make antimicrobial copper an ideal candidate for surface protection, not just against Covid-19, but the common cold, the flu, multi-resistant bacteria, and future pandemics. Studies have proven the cost-effectiveness of copper in the case of HAI.^21^ Using a thin sheet of copper, as in the copper sticker tested in this study, to cover surfaces may further reduce cost of material and improve feasibility of use on an industrial scale. It has been shown to be highly antibacterial and practical in a University dorm setting.^22^

## Conclusion

Copper is an effective and practical means of protection against fomite transmission of SARS-CoV-2.

## ACKNOWLEDMENTS

We thank Dr Marcela Ferres and all her team at Infectology Lab for provide positive patient’s sample to isolate infectious SARS-CoV-2. We also thank to Dr Rafael Medina for kindly provide basic protocol of Influenza’s plaque assay which was modified for SAR-CoV-2 plaque assay. We also thank to Silvia Varietti and Stephanie Mahuzier from Diagnochile SpA for kindly provide SARS-CoV-2 antigen test used in this study.

## References

1. Hashana MR, Smoll N, King K, et al. Epidemiology and clinical features of COVID-19 outbreaks in aged care facilities: A systematic review and meta-analysis. E Clin Med. Volume 33, 100771, March 01, 2021. https://doi.org/10.1016/j.eclinm.2021.100771.

2. Ioannidis JPA. Global perspective of COVID-19 epidemiology for a full-cycle pandemic. Eur J Clin Invest. 2020;00:e13423. https://doi.org/10.1111/eci.13423.

3. Campos EV, Pereira AE, Oliveira JL, et al. How can nanotechnology help to combat COVID-19? Opportunities and urgent need. J Nanobiotechnol (2020) 18:125. https://doi.org/10.1186/s12951-020-00685-4.

4. Kchaou M, Abuhasel K, Khadr K, et al. Surface Disinfection to Protect against Microorganisms: Overview of Traditional Methods and Issues of Emergent Nanotechnologies. Appl. Sci.2020,10, 6040; doi:10.3390/app10176040.

5. Hutasoita N, Kennedyb B, Hamilton S, et al. Sars-CoV-2 (COVID-19) inactivation capability of copper-coated touch surface fabricated by cold-spray technology. Manufacturing Letters 25 (2020) 93s–97, DOI: 10.1016/j.mfglet.2020.08.007

6. Cortesa AA, Zuñiga JM. The use of copper to help prevent transmission of SARS-coronavirus and influenza viruses. A general review. Diagn Microbiol Infect Dis. 2020 Dec; 98(4): 115176. doi: 10.1016/j.diagmicrobio.2020.115176.

7. Grass G, Rensing C, Solioz M. Metallic Copper as an Antimicrobial Surface. Appa Environ Microbiol. 2011, Vol. 77(5),1541–1547. doi:10.1128/AEM.02766-10.

8. https://www.epa.gov/newsreleases/epa-registers-copper-surfaces-residual-use-against-coronavirus

9. Doremalen NV, Morris DH, Holbrook MG, et al. Aerosol and Surface Stability of SARS-CoV-2 as Compared with SARS-CoV-1. N Eng J Med 2020, 382;16. DOI: 10.1056/NEJMc2004973.

10. Warnes SL, Little ZR, Keevil CW. Human Coronavirus 229E Remains Infectious on Common Touch Surface Materials. mBio. November/December 2015, 6(6)e01697–15. doi: 10.1128/mBio.01697-15, doi: 10.1128/mBio.01697-15.

11. Espirito Santo, C., P. V. Morais, and G. Grass. 2010. Isolation and characterization of bacteria resistant to metallic copper surfaces. Appl. Environ. Microbiol. 76:1341–1348. doi: 10.1128/AEM.01952-09.

12. Quaranta, D., et al. 2011.Mechanisms of contact-mediated killing of yeast cells on dry metallic copper surfaces.Appl. Environ. Microbiol.77:416–426,doi:10.1128/AEM.01704-10

13. Wheeldon, L. J., et al. 2008. Antimicrobial efficacy of copper surfaces against spores and vegetative cells of Clostridium difficile: the germination theory. J. Antimicrob. Chemother. 62:522–525, doi: 10.1093/jac/dkn219

14. Warnes SL, Keevil CW. Mechanism of Copper Surface Toxicity in Vancomycin-Resistant Enterococci following Wet or Dry Surface Contact. App Envir Micro 2011,77(17);6049–59, doi: 10.1128/AEM.00597-11.

15. Molteni C, Abicht HK, Solioz M. Killing of Bacteria by Copper Surfaces Involves Dissolved Copper. App Envir Micro 2011,76(12);4099–4101, doi: 10.1128/AEM.00424-10.

16. Warnes SL, Keevil CW (2013) Inactivation of Norovirus on Dry Copper Alloy Surfaces. PLoS ONE 8(9): e75017. doi:10.1371/journal.pone.0075017.

17. Bryant C, Wilks SA, Keevil CW. Rapid inactivation of SARS-CoV-2 on copper touch surfaces determined using a cell culture infectivity assay. bioRxiv preprint doi: https://doi.org/10.1101/2021.01.02.424974.

18. Xie C, Zhao H, Li K, et al. The evidence of indirect transmission of SARS-CoV-2 reported in Guangzhou, China. BMC Public Health (2020) 20:1202. https://doi.org/10.1186/s12889-020-09296-y.

19. Lessells R, Moosa Y, de Oliveira T. Report into a nosocomial outbreak of coronavirus disease 2019 (COVID-19) at Netcare St. Augustine’s Hospital. KwaZulu1Natal Research Innovation and Sequencing Platform (KRISP). https://www.krisp.org.za/news.php?id=421(pdf) 2020

20. Salgado CD, Sepkowitz KA, John JF, et al. Copper Surfaces Reduce the Rate of Healthcare-Acquired Infections in the Intensive Care Unit. Infect Control Hosp Epidemiol. 2013;34(5):479–486, doi: 10.1086/670207.

21. Michels HT, Keevil CW, Salgado CD, et al. From Laboratory Research to a Clinical Trial: Copper Alloy Surfaces Kill Bacteria and Reduce Hospital-Acquired Infections. Health Environ Research Design J. 2015;9(1):64–79, doi: 10.1177/1937586715592650.

22. Lu K, Mendez N. Antimicrobial Copper Foil Reduces Bacterial Contamination and Load on Door Handles of Loyola Marymount University Dormitory. Biomed J Sci & Tech Res. 2019;21(5):16179–16182. DOI: 10.26717/BJSTR.2019.21.003663.

